# Identification of Mitosis Stages Using Artificial Neural Networks for 3D Time Lapse Cell Sequences

**DOI:** 10.1101/2024.02.12.579090

**Authors:** Tolga Dincer, Johannes Stegmaier, Abin Jose

## Abstract

Cells, the fundamental units of life, are central to medical research, particularly in cancer studies due to their rapid, uncontrolled division. Understanding cell behavior is crucial, with a focus on mitosis, which has distinct cell division stages. However, precise detection of these phases, especially mitosis initiation in 3D, remains an underexplored research area. Our work explores 3D cell behavior, leveraging the increasing computational capabilities and prevalence of 3D imaging techniques. We introduce diverse 3D Convolutional Neural Network (CNN) architectures such as a base 3D CNN model, 3D CNN binary model, and 3D CNN pairwise model. An ensemble model based on the 3D CNN architectures shows higher classification accuracy on two time-series datasets. This research gives better insights into understanding cell behaviour in a multidimensional manner, contributing to medical research. To the best of our understanding, we are the first to delve into the utilization of Convolutional Neural Network architectures for the 3D classification of mitosis stages.

## 1. INTRODUCTION

In the field of medical research, cells, the fundamental units of life, hold a pivotal position, especially in the context of cancer studies. Cell division, a fundamental biological process for development, growth, and tissue repair, involves a single mother cell giving rise to two daughter cells [1]. The urgency of comprehensively understanding cell behavior is underscored by diseases like cancer, characterized by rapid and uncontrolled cell growth and division [1]. Mitosis, the most common form of cell division, consists of distinct phases [2]. While recent research has made notable progress in cell tracking and segmentation [3], specifically cell-cycle detection, particularly the initiation of mitosis, remains relatively unexplored. Advances in image processing [4, 5] and deep learning [6, 7] have revitalized the pursuit of automated cell-cycle detection [8]. Notably, cell-cycle detection and classification research has predominantly focused on 2D sequences [9, 10], overlooking the untapped potential of insights that 3D imaging techniques can provide.

With increasing computational power and the emergence of 3D imaging, the exploration of cell behavior in 3D videos has become a compelling avenue for investigation, forming the core of this paper. In this work, we present multiple methods for automatic cell-cycle classification tailored to 3D + time (3D+t) data. Our primary objective is to categorize frames of 3D video sequences into interphase (pre-mitosis), mitosis, and post-mitosis phases in an end-to-end manner, as defined in [9].

Our focus extends to identifying the precise frame where a cell initiates its mitosis cycle and determining the duration of mitosis until it divides into two daughter cells. While classifying frames into specific mitosis phases has been explored for 2D sequences [8], our research focuses on relatively uncharted territory of 3D sequences. To evaluate our algorithms, we use bounding boxes to track individual cells over time. Leveraging deep learning techniques, particularly various 3D CNN [11] architectures, we explore the classification of cell-cycle stages of 3D+t data.

### 1.1. Related work

As this paper focuses on detecting cell mitosis frames in microscopy images, we explored 3D models in various data types and tasks. For example, Singh et al. [12] presented models for segmentation and classification using 3D data, such as MRI, fMRI, and PET scans, often employing 3D CNN [11] architectures. Although limited in 3D video research, existing methods use 3D CNNs and RNN-based models [13]. For instance, Tran et al. [11] introduced 3D CNNs for spatio-temporal feature extraction from video sequences, and CNN + LSTM [14, 15] and two-stream [16] architectures are common for video classification.

In the context of cell-cycle stage identification for 2D data, LiveCellMiner [8], an open-source tool for 2D+t microscopy image data analysis, provides feature extraction, mitosis stage identification, cell segmentation, and tracking. It utilizes classical features [17], LSTM networks [18, 19], and CNN architectures. Additionally, Jose et al. [9] proposed RNN-based architectures, combining convGRU [20] layers with CNNs for 2D + time cell-cycle stage classification. For 3D cell-cycle stage identification, Du et al. [21] manually extracted 3D features and used SVM [22] classification. To the best of our knowledge, there’s currently a gap in deep learning-based frame-wise classification models for cell-cycle stages for 3D+t data.

### 1.2. This paper

Our paper addresses the gap in frame-wise classification for cell-cycle stages for 3D+t sequences, drawing inspiration from related works in medical imaging, video classification, and cell-cycle stage identification [11, 23, 24, 9]. We propose 3 main models and an ensemble model in this paper such as, 1) Frame-wise classification of cell-cycle stages using 3D CNN architecture. 2) Classification of cell state transition and non-transition frames using a consecutive frames (pair-wise) approach. 3) 3D CNN model to specifically distinguish between mitosis and non-mitosis frames. 4) Combination of previous three models to create an ensemble model that improves upon the base 3D CNN approach. Experiments were conducted on Fluo-N3DH-SIM+ dataset from Cell Tracking Challenge and we also created another synthetic dataset for experimental validation. The ensemble model showed higher classification performance.

### 1.3. Organization of the paper

The paper is organised as follows: Section 2 introduces the proposed approach. Training details are elaborated in Section 3. Experimental results are discussed in Section 4. Concluding remarks are discussed in Section 5.

## 2. PROPOSED APPROACH

### 2.1. Base 3D CNN model

Our primary objective is to classify video frames into three classes: S/G2 (pre-mitosis), M (mitosis), and G1 (postmitosis), numerically encoded as 0, 1, and 2. We propose a 3D CNN architecture (Fig. 1), inspired by [25], which takes each frame as input for classification. This approach focuses solely on spatial features, excluding temporal features.

**Fig. 1:**
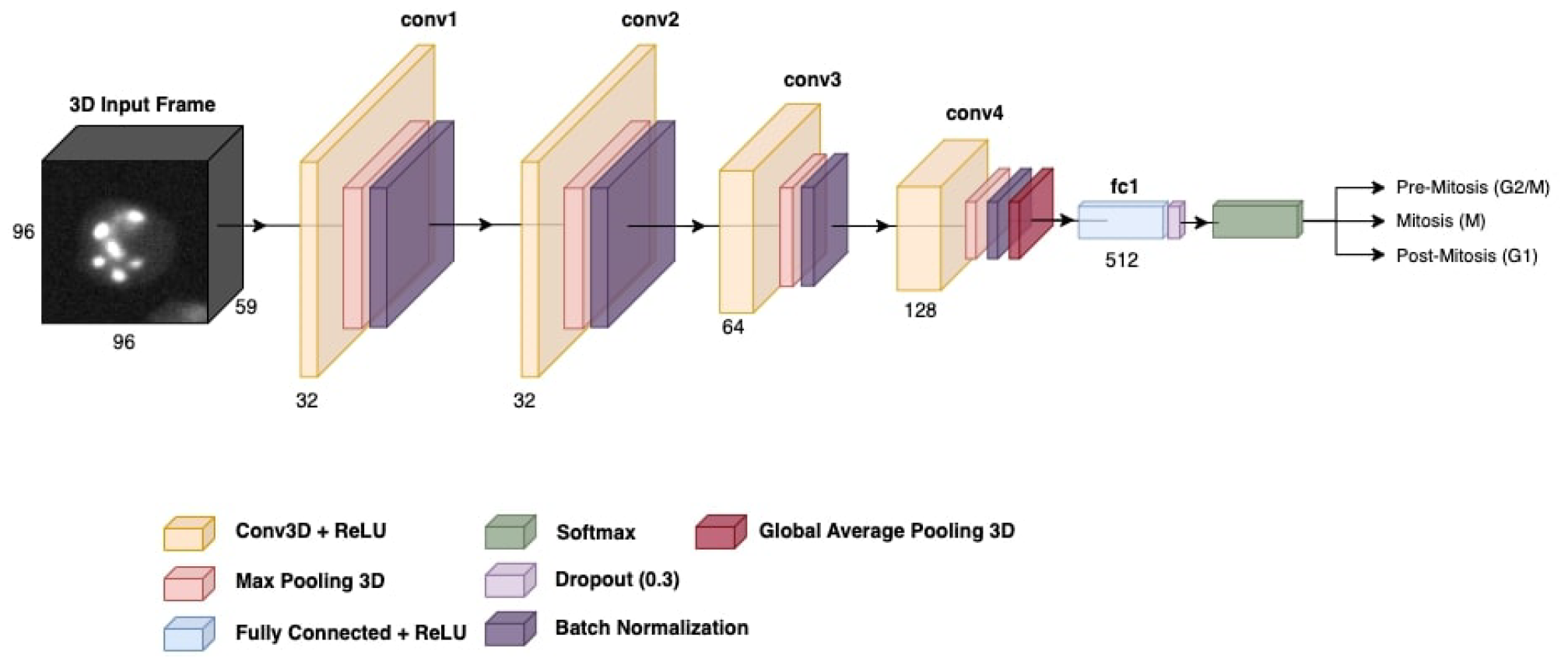
3D CNN model, showing one frame of the 3D cell splitting sequence. The softmax layer predicts the cell state and classifies it into pre-mitosis, mitosis or post-mitosis.

Our base 3D CNN model processes individual frames with (width, height, depth) dimensions. It begins with a convolutional layer (Conv3D) consisting of 32 filters and a kernel size of (3 × 3 × 3). After convolution, we apply Max-Pooling3D to reduce dimensionality. All models in this paper employ a (2 × 2 × 2) pooling size. We introduce batch normalization (BN) after max pooling. The second conv3D layer mirrors the first one with 32 filters, max pooling, and BN. The third and fourth layers utilize 64 and 128 filters, followed by max pooling and BN. After the final BN layer, global average pooling is applied, addressing computational limitations. A fully connected layer with 512 parameters is introduced after global average pooling, followed by a dropout with a 0.3 probability.

### 2.2. 3D CNN binary model

We introduce a binary 3D CNN model to specifically differentiate between mitosis and non-mitosis frames. The only modification done to the base 3D CNN model is the use of a sigmoid layer instead of a softmax layer for the classification output as shown in Fig.2 and a corresponding change in the loss function. The sigmoid model enables the network to predict the final state of the model as either mitosis or non-mitosis.

**Fig. 2:**
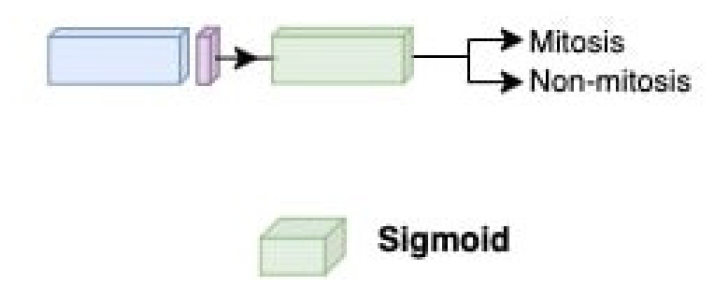
Binary classification model, with sigmoid layer at the end which predicts the state of the cell as mitosis or non-mitosis.

### 2.3. 3D CNN pairwise model

To enhance the base 3D CNN model, we introduce a pairwise frames approach where Conv3D layers apply identical convolutions to two channels containing the current and the next frame, enabling the incorporation of spatial feature evolution between the two frames. The model learns to map a concatenated frame pair to stage transition codes in the form of 00, 01, 11, 12, 22, 20. The 01 and 12 classes represent transitions into and out of the mitosis stage. Data points are shaped as (height × width × depth × 2), with the channel dimension set to 2. This concept draws inspiration from [23], which applies a similar strategy to binary classification of 4D fMRI data. Our architecture maintains the same layers as previous models in this paper, with the key change being the output layer, which is transformed into a six-class softmax. For a visual summary of the modifications to the original base 3D CNN model given in Fig. 1, refer to Fig.3.

**Fig. 3:**
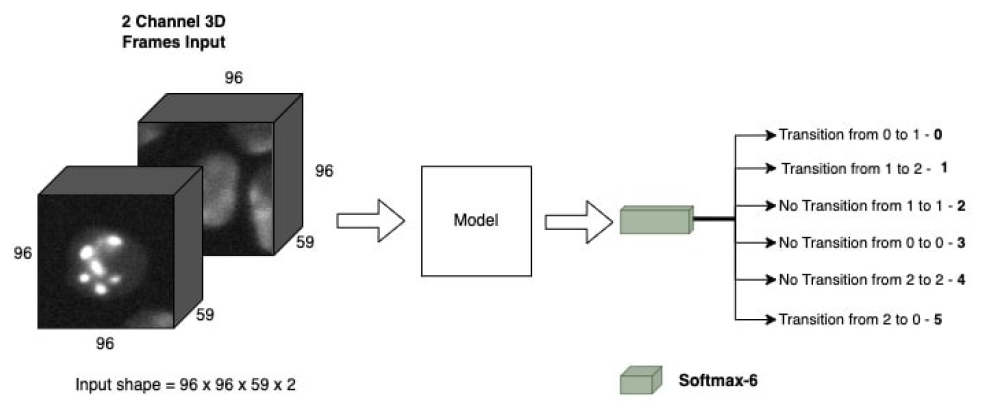
Transition frames classification model, predicting the transition states.

### 2.4. Ensemble model

In order to improve the results of the 3D CNN base model we employ an ensemble model that combines results from pair-wise and binary model with the base 3D CNN model for three class classification.

### 2.5. Voting for ensemble model

Voting-based ensemble methods, such as majority voting [26], combine predictions from multiple base learners to determine the final prediction. Each base learner independently predicts a class label, and the final prediction is based on the majority vote among these individual predictions. Base 3D CNN model naturally vote for their current frame prediction. 3D CNN Binary model, on the other hand, have two options: they can vote for mitosis (1), or they can vote for both 0 (pre-mitosis) and 2 (post-mitosis). The reason for this dual voting in binary model is to ensure that the voting outcome remains consistent even when the base 3D CNN model predicts either 0 or 2. By voting for both labels (0 and 2), these labels receive additional votes, which helps maintain the majority, regardless of whether the base 3D CNN model predicts 0 or 2. In the 3D CNN pairwise model, predictions are based on the labeling of both the current and next frames. For instance, a prediction of 0 corresponds to voting for 0 in the current frame and 1 in the next frame (see Fig. 3 for class encoding).

## 3. TRAINING DETAILS

### 3.1. Training the base 3D CNN model

Normalization is applied to each sample before data augmentation. For all of our 3D CNN models, rotation and random flipping were applied. Rotation is applied based on a random selection of a plane (xz, xy, and yz). Afterwards, a random value is selected from a list of angles to apply rotation to the 3D image along the initially chosen plane. The angles are: [-20, -10, -5, 5, 10, 20]. For augmentation via flipping, the image is flipped along one or multiple axes. Each flip along an axis has a probability of 0.5. While traditional augmentation seeks to increase the data used in training, due to the computation cost of using many 3D data samples, here the augmentations are applied to the training data randomly in each epoch. This way the models see variants of the data and generalize better.

The loss function used in this classification task is sparse categorical cross-entropy. Sparse categorical cross-entropy only differs from normal categorical cross-entropy in that the labels are not one-hot encoded, such as 00, 01, 10, but are given as integers, such as 0, 1, and 2. The cross-entropy loss function used is:

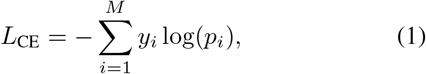

where *p*_*i*_ represents the softmax probability for the *i*^*th*^ class, and *y*_*i*_ represents the true label [27]. To initiate training, a learning rate of 0.00018 and 0.0002 is used for the Fluo-N3DH-SIM+ and the simulated dataset respectively and a batch size of two was used for all the models’ training.

### 3.2. Training the binary model and pairwise model

For binary classification, binary cross-entropy is used as the loss function. The pre-mitosis and post-mitosis classes are merged into the non-mitosis class. For the six-class pairwise model, the dataset is modified in such a way that each data point includes consecutive frames that forms the two channels of the input. The learning rate for the binary model is defined as 0.00018 and 0.00016 for Fluo-N3DH-SIM+ and the simulated dataset respectively. For the pairwise model, these learning rates are 0.0001 and 0.00016 respectively. The rest of the training parameters are the same as before with the loss function defined above.

## 4. EXPERIMENTAL SETUP

### 4.1. The 3-class dataset

To perform the classification tasks described earlier, we utilize the **Fluo-N3DH-SIM+ dataset** from the Cell Tracking Challenge [28]. This dataset was chosen for its high quality and resolution, featuring easily identifiable mitosis frames. Fluo-N3DH-SIM+ is an artificial dataset generated using MitoGen [29], serving as a benchmark for the Cell Tracking Challenge. MitoGen produces 3D time-lapse sequences of synthetic fluorescence-stained cell populations with ground truth. In the original dataset, multiple cells move through various cell stages over time.

In this paper, we extract individual cell bounding boxes by leveraging tracking and mask ground truth by using the z-axis slice with the largest mask for extracting bounding boxes and individual cell sequences. Thus, daughter cells following mitosis are tracked as separate videos. However, the sequences share frames between each other. For example, 2 daughter cells forming as a result of a mitosis event each has their own sequence but they share the frames before mitosis. These frames are eliminated to avoid duplicate data during training.

In Fig. 4, mitosis frames are distinguishable by higher chromosomal intensities. Labeling challenges primarily involve differentiating between G1 (post-mitosis) and G2/S (pre-mitosis) phases. Our labeling criteria entail marking frames as G1 when cells begin growing after mitosis until reaching a stable size. Subsequently, we label frames as G2/S until the cell enters the mitosis phase. Each bounding box in this dataset has dimensions (96, 96, 59), and each sequence comprises 150 frames. The dataset contains a total of 40 sequences.

**Fig. 4:**
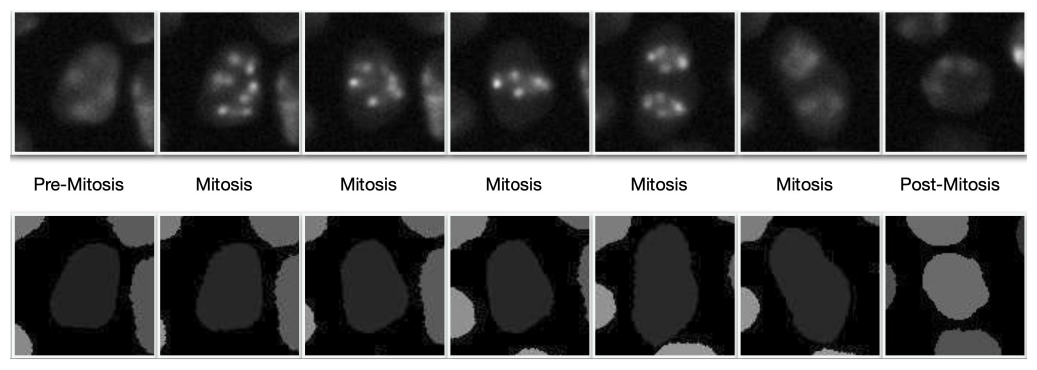
Example cell sequence in different stages of mitosis in first row, with respective labels in the middle row and mask in the bottom row. The z-axis slice with the largest masks are visualized.

### 4.2. The 3-class simulated dataset

To further validate the performance of the model, we simulated a second dataset in the same manner as the first one. The dataset simulator by Masaryk University [30] was used for simulation. Each bounding box in this dataset has dimensions (190, 190, 60) and each sequence is 100 frames long. There are a total of 34 sequences in this dataset. As before, the duplicate frames are omitted. The magnification parameter used for the simulation is Zeiss Plan-Apochromat 100*x/*1.40 Oil, the simulated microscope is Zeiss 200M and the confocal unit is Yokogawa CSU-10. The simulated camera is CoolSNAP HQ. The excitation filter and the emission filter are 640*/*6 Andor Laser and 685*/*40 Cy5 Semrock consecutively.

### 4.3. Evaluation criteria and hyperparameters

To evaluate for each dataset, we reserve one full sequence of 150 and 100 frames respectively for testing. The training set is split into training and validation using a 80/20 percent split. We use accuracy, precision, recall and F1-score as our evaluation metrics, which are calculated for each of the classes. In our opinion, the most important evaluation criteria are the F1-score for the mitosis class and the overall accuracy.

## 5. EXPERIMENTAL RESULTS

### 5.1. Results for Fluo-N3DH-SIM+ dataset

From the confusion matrix results shown in Fig. 5(a), it is evident that the base 3D CNN model performs well in classifying mitosis frames, achieving 100% precision and 90% recall for the mitosis class. However, there is a notable bias towards pre-mitosis (0), with almost half of the post-mitosis (2) frames being misclassified as pre-mitosis (0). The base 3D CNN model attains an 82% accuracy rate with an F1-score of 0.947 for the mitosis class, while the F1-score for post-mitosis is comparatively lower at 0.658.

**Fig. 5:**
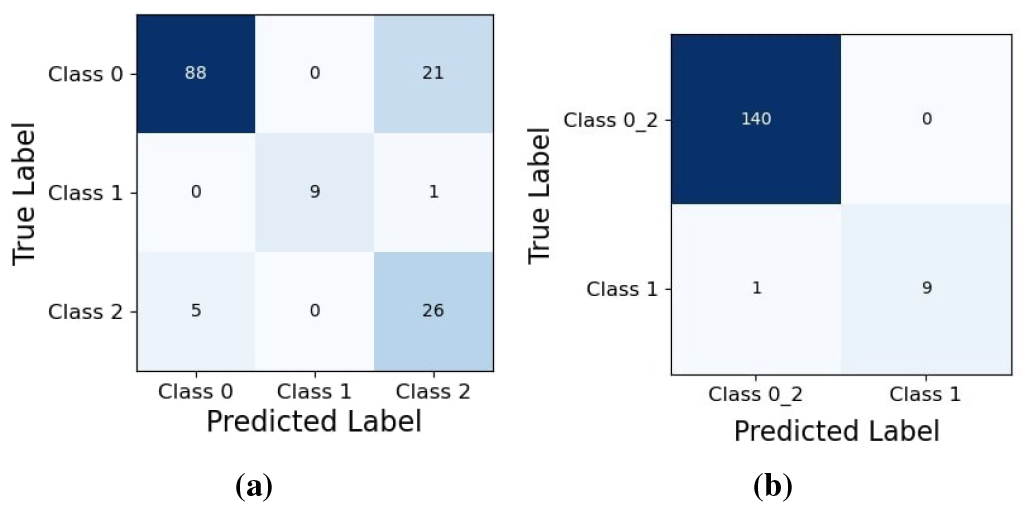
Confusion matrices for (a) three-class classification and (b) binary models for Fluo-N3DH-SIM+ dataset.

A detailed summary of the evaluation metrics for the models are provided in Tables 1 and Table 2. The confusion matrix for the binary model in Fig. 5(b) shows only one false classification for mitosis frames. The confusion matrix for the pairwise model is shown in Fig. 6(a) and the valuation metrics are provided in Table 3 and Table 4. Combining the base 3D CNN, binary and the pairwise model, the ensemble model’s results, as shown in Fig. 6(b) and Table 1 and Table 2, improve upon the base 3D CNN in almost all metrics. The ensemble model enhances accuracy to approximately 92.667%, demonstrating an approximate 10% increase over the base 3D CNN model. Notably, the F1-score for both pre- and post-mitosis classes experience significant improvements, while the F1-score for the mitosis class remains consistent.

**Table 1:** Results for Fluo-N3DH-SIM+ - Accuracy and Precision in %.

| Model | Accuracy | Precision |  |  |
| --- | --- | --- | --- | --- |
|  |  | 0 - Pre-Mitosis | 1 - Mitosis | 2 - Post-Mitosis |
| Base 3D CNN | 82.000 | 94.624 | 100.00 | 54.167 |
| Ensemble | **92.667** | **96.262** | **100.00** | **79.412** |

**Table 2:**
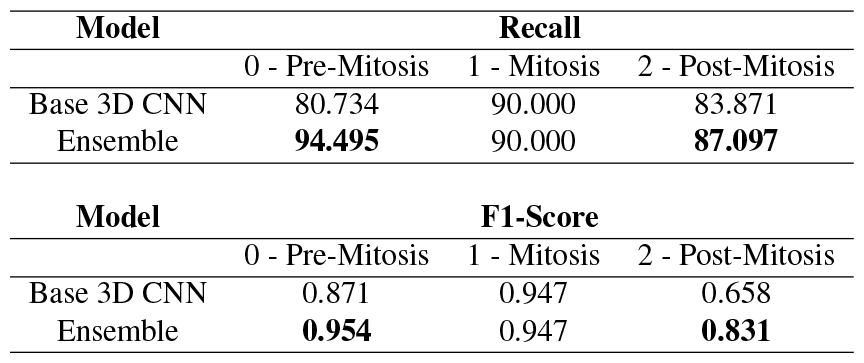
Results for Fluo-N3DH-SIM+ - Recall in % and F1-scores.

**Table 3:** Fluo-N3DH-SIM+ pairwise model results - Accuracy and Precision in %.

| Model | Accuracy | Precision |  |  |  |  |  |
| --- | --- | --- | --- | --- | --- | --- | --- |
|  |  | 01 | 12 | 11 | 00 | 22 | 20 |
| Pairwise Model | 77.852 | 100.00 | 100.00 | 100.00 | 94.253 | 46.154 | None |

**Table 4:** Fluo-N3DH-SIM+ pairwise model results - Recall in % and F1 scores.

| Model | Recall |  |  |  |  |  | F1 Score |  |  |  |  |  |
| --- | --- | --- | --- | --- | --- | --- | --- | --- | --- | --- | --- | --- |
|  | 01 | 12 | 11 | 00 | 22 | 20 | 01 | 12 | 11 | 00 | 22 | 20 |
| Pairwise model | 100.00 | 50.000 | 87.500 | 77.358 | 82.759 | 0.000 | 1.000 | 0.667 | 0.933 | 0.849 | 0.593 | None |

**Fig. 6:**
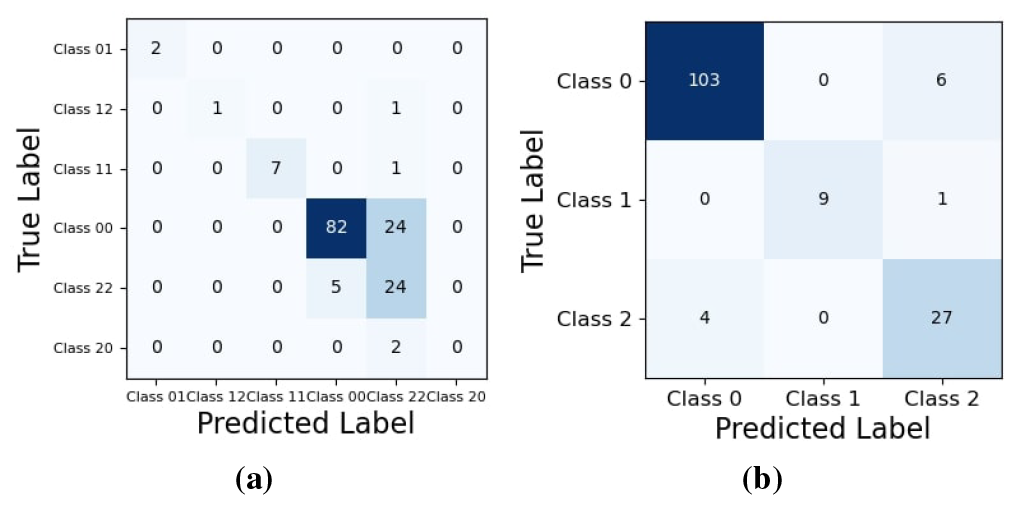
Confusion matrices for (a) pairwise model, and (b) ensemble model for Fluo-N3DH-SIM+ dataset.

### 5.2. Results for the simulated dataset

The simulated dataset, constrained by GPU limitations due to larger bounding boxes, employs a smaller training dataset. The base 3D CNN model, as shown in Fig. 7(a), exhibit significantly improved differentiation between pre- and post-mitosis classes. The binary model misclassifies only one frame of mitosis as shown in Fig. 7(b). The confusion matrix for the pairwise model, as shown in Fig. 8(a) shows much better differentiation of the frames without any cell-stage transitions compared to the previous dataset. The metrics for the pairwise model are summarised in Tables 7 and 8. As with the previous dataset, the ensemble model surpasses the base 3D CNN model in almost every metric.

**Table 5:** Results for the simulated dataset - Accuracy and Precision in %.

| Model | Accuracy | Precision |  |  |
| --- | --- | --- | --- | --- |
|  |  | 0 - Pre-Mitosis | 1 - Mitosis | 2 - Post-Mitosis |
| Base 3D CNN | 94.000 | 92.188 | 88.889 | 100.00 |
| Ensemble | 96.907 | 95.161 | 100.00 | 100.00 |

**Table 6:** Results for the simulated dataset - Recall in % and F1-scores.

| Model | Recall |  |  |
| --- | --- | --- | --- |
|  | 0 - Pre-Mitosis | 1 - Mitosis | 2 - Post-Mitosis |
| Base 3D CNN | 100.00 | 88.889 | 84.375 |
| Ensemble | 100.00 | 88.889 | 93.103 |

**Table 6:** Results for the simulated dataset - Recall in % and F1-scores.
| Model | F1-Score |  |  |
| --- | --- | --- | --- |
|  | 0 - Pre-Mitosis | 1 - Mitosis | 2 - Post-Mitosis |
| Base 3D CNN | 0.959 | 0.889 | 0.915 |
| Ensemble | 0.975 | 0.941 | 0.964 |

**Table 7:**
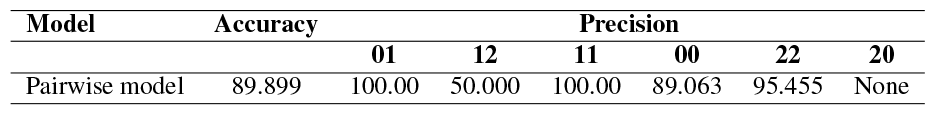
Simulated dataset pairwise model results - Accuracy and Precision in %.

**Table 8:** Simulated dataset pairwise model results - Recall in % and F1 scores.

| Model | Recall |  |  |  |  |  | F1 Score |  |  |  |  |  |
| --- | --- | --- | --- | --- | --- | --- | --- | --- | --- | --- | --- | --- |
|  | 01 | 12 | 11 | 00 | 22 | 20 | 01 | 12 | 11 | 00 | 22 | 20 |
| Pairwise Model | 100.00 | 100.00 | 100.00 | 100.00 | 70.000 | 0.000 | 1.000 | 0.667 | 1.000 | 0.942 | 0.808 | None |

**Fig. 7:**
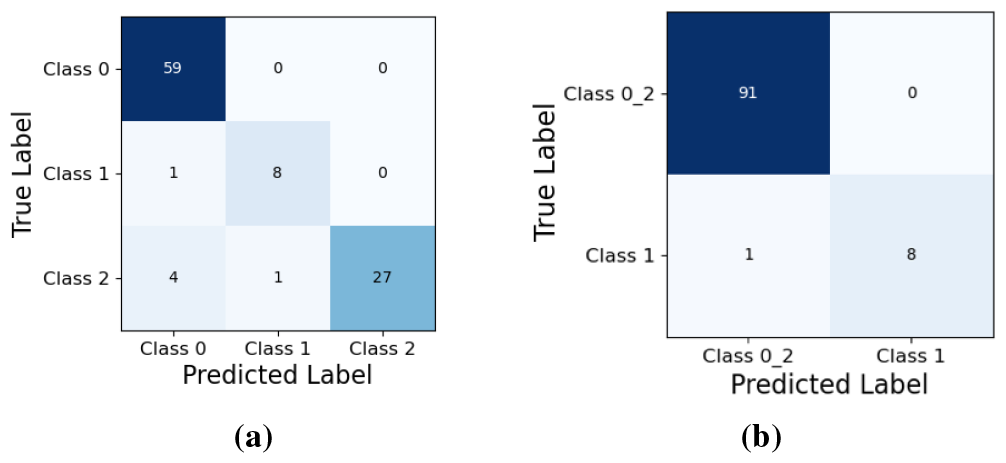
Confusion matrices for (a) three-class classification and (b) binary model for simulated dataset.

**Fig. 8:**
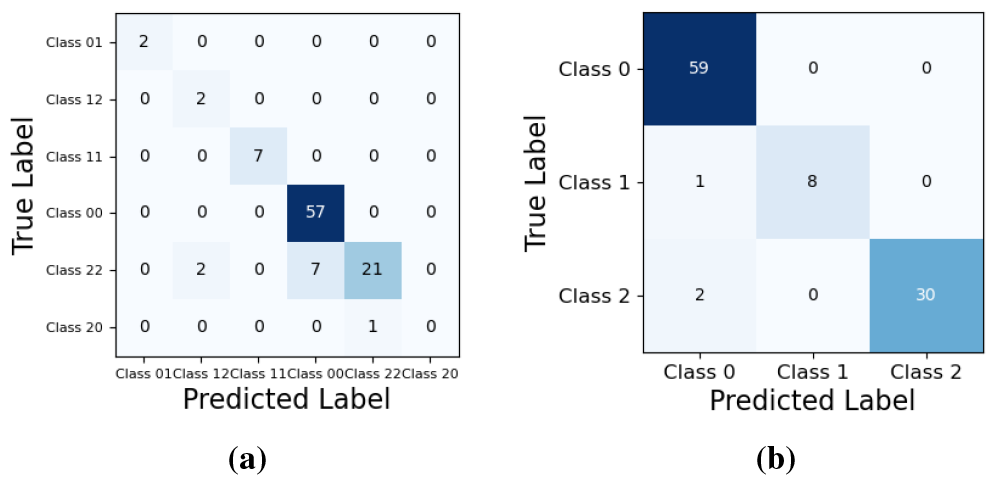
Confusion matrix for (a) pairwise and (b) ensemble model for simulated dataset.

The confusion matrix for the ensemble model, as shown in Fig. 8(b), differentiates between pre- and post-mitosis classes better than the base model. The ensemble model also provides an improvement of F1-scores of the post-mitosis (2) class provided in Table 6. Comparing the results of both the base 3D CNN model and the ensemble model in Tables 5 and 6, it becomes clear that the ensemble model provides an improvement on the base 3D CNN model, achieving an F1-score of 0.941 for the mitosis class and the best F1-scores for all classes overall. The ensemble model achieves 96.907% accuracy surpassing the base model by 2.907%. For the base 3D CNN model, we recommend a GPU with RAM between 20 and 24 gigabytes (GB) (TITAN RTX in our case).

## 6. CONCLUSIONS AND FUTURE WORK

3D+t data offers a comprehensive perspective for studying cell division, combining spatial and temporal features to provide insights into cellular structures. This depth and spatial context are essential for understanding cell shapes and characteristics, surpassing the limitations of 2D videos. Given the novelty of our task and the absence of existing deep learning models for frame-wise cell-cycle stage classification, we explored and compared two different models. The ensemble model, combining the base 3D CNN model with the binary model and the pairwise frames model, outperforms the base 3D CNN model. Utilizing deeper architectures could enhance results; during our architectural exploration, we observed that deeper and more complex models outperformed simpler ones. However, computational costs for deep learning on 3D+t data must be considered. With increased computational power, we anticipate that more complex models will yield improved results. As future work, we want to explore the problem of tracking daughter cells after cell division for 3D sequences. Leveraging temporal features and using transformers with long-range temporal dependencies and unsupervised training strategies are other possible research directions.

